# Electrosensory and metabolic responses of weakly electric fish to changing water conductivity

**DOI:** 10.1101/2023.10.18.562945

**Authors:** Shannon D. Wiser, Michael R. Markham

## Abstract

Weakly electric Gymnotiform fishes use self-generated electric organ discharges (EODs) to navigate and communicate. The electrosensory range for these processes is a function of EOD amplitude, determined by the fish’s electric organ (EO) output and the electrical conductivity of the surrounding water. Anthropogenic activity, such as deforestation, dams, and industrial/agricultural runoff, are known to increase water conductivity in neotropical habitats, likely reducing the electrosensory range of these fish. We investigated whether fish modulate EO output as means of re-expanding electrosensory range after a rapid increase in water conductivity in the pulse-type *Brachyhypopomus gauderio* and the wave-type *Eigenmannia virescens*. Furthermore, because EOD production incurs significant metabolic costs, we assessed whether such compensation is associated with an increase in metabolic rate. Following the conductivity increase *B. gauderio* increased EOD amplitude by 20.2 ± 4.3% over six days but with no associated increase in metabolic rate, whereas the EOD amplitude of *E. virescens* remained constant, accompanied by an unexpected decrease in metabolic rate. Our results suggest that *B. gauderio* uses a compensation mechanism that requires no metabolic investment, such as impedance matching, or a physiological tradeoff wherein energy is diverted from other physiological processes to increase EO output. These divergent responses between species could be the result of differences in reproductive life history or evolutionary adaptations to different aquatic habitats. Continued investigation of electrosensory responses to changing water conditions will be essential for understanding the effects of anthropogenic disturbances on gymnotiforms, and potential physiological mechanisms for adapting to a rapidly changing aquatic environment.

## Introduction

Anthropogenic activity such as deforestation, dams, mining, industrial waste disposal, and agricultural runoff negatively impact the estimated 5000 species of fishes living in the freshwater ecosystems of South America (Reis et al., 2016) by degrading water quality, crating barriers that inhibit aquatic migration, and altering seasonal weather patterns (Castello & Macedo, 2016, Coe et al., 2011; Costa & Foley, 1997). Changes in ionic concentrations, especially salinity, are a common consequence of many anthropogenic disturbances, resulting in corresponding changes in the water’s electrical conductivity. Although most freshwater fishes can adapt to changes in water conductivity, experienced as osmotic stress (Kultz, 2015), the South American weakly electric gymnotiform fishes may experience an additional type of stress due to the potential effects of water conductivity on their electric signals.

Gymnotiform fishes generate electric fields, known as electric organ discharges (EODs), and detect distortions of these electric fields for sensory processes such as navigation, communication, and foraging. In all gymnotiform species, the EOD is produced by an electric organ (EO) that extends bilaterally along the body and into the tail. The EO is comprised of electrogenic cells, known as electrocytes, that produce the EOD via coordinated action potentials. Electric current, generated by ionic currents through the electrocyte voltage-gated ion channels, is directed along the fish’s body and produces an electrical current that flows through the surrounding water (Markham, 2013). Gymnotiform species are categorized either as wave-type fishes that produce EODs at continuous frequencies of ∼100-2000 Hz, or as pulse-type fishes that generate EODs at low intermittent rates of ∼10-100 Hz. In all species examined to date, EOD production incurs significant metabolic costs: as high as 30% of the daily energy budget in wave fishes and 20% in pulse fishes. (Lewis et al., 2014; Salazar & Stoddard, 2008; Salazar et al., 2013).

EOD amplitude (EODa; measured as electrical potential in water) likely coincides with the fish’s electrosensory range – the distance at which fish can detect objects or communicate with conspecifics (Rasnow, 1996). In accordance with Ohm’s Law, which states that for a constant electrical current, electrical potential is inversely related to conductivity, EODa is determined by the fish’s electric organ (EO) output and water conductivity. Accordingly, increased conductivity reduces EODa if EO output is constant, while decreased conductivity increases EODa at a constant EO output. In a shift from low to high water conductivity, some species have shown decreased sensory performance (MacIver et al., 2001), suggesting that increased water conductivity may degrade a fish’s ability to forage or detect prey.

Many species of electric fish modulate EO output over minutes to hours in response to circadian cues and social encounters (Markham et al., 2009; Markham & Stoddard, 2005), which suggests a potential mechanism to compensate for the effects of increased water conductivity on EODa by increasing EO output accordingly. This led us to investigate whether fish increase EODa to compensate for the effects of increased water conductivity. If fish do compensate by increasing EO output, an important second question that follows is whether the increase in EO output requires additional metabolic investment in EOD production. Given the high metabolic costs of EOD production under normal circumstances, any additional metabolic investment could make EOD production physiologically unsustainable forcing a decrease in sensory performance, or a metabolic tradeoff with physiological functions such as immunity and reproduction.

To address these questions, we studied the electrosensory and metabolic responses of pulse- and wave-type gymnotiform species to rapid increases in water conductivity. This comparative approach allowed us to evaluate whether species with different EOD rates and with different life histories might exhibit different response strategies. We measured EODa as a metric of electrosensory performance and we measured metabolic responses by intermittent flow respirometry. In increased water conductivity, we hypothesized that if the fish increases EO output to compensate for the reduced EODa, this response will cause an associated increase in metabolic rate.

Conversely if the fish does not increase EO output, it will maintain a constant metabolic rate, likely forcing a tradeoff in electrosensory performance.

## Materials and Methods

### Animals and water treatment

The fish used for this study were the wave-type gymnotiform *Eigenmannia virescens* and the pulse-type gymnotiform *Brachyhypopomus gauderio. E. virescens* were obtained through tropical fish importers and *B. gauderio* were obtained from an on-campus breeding colony at the University of Oklahoma. Fish were housed in an indoor recirculating aquarium system with a 12L:12D light cycle, at 26°C, and fed live oligochaete black worms *ad libitum*. Water for the animals was prepared from reverse-osmosis purified and deionized water. Water conductivity was controlled using a concentrated pH-buffered saline solution (Walter’s Solution) consisting of deionized water containing (in mM): CaSO_4_•2H_2_0 (732), MgSO_4_ (83), KCl (107), NaH_2_PO_4_•H_2_0 (17), FeC_6_H_5_O_7_ (7). Fish care and experimental protocols were approved by the Institutional Animal Care and Use Committee of the University of Oklahoma.

### Experimental Timeline

EOD recordings and respirometry experiments were performed separately but under standardized water conductivity parameters. Fish were acclimated to the Low Conductivity condition (100 ± 50 μS) for a minimum of 7 days before data collection began to eliminate stress related to changing water conditions during baseline measurements. Measurements were collected within the Low Conductivity condition for two days (Day -2 and Day -1) before Walter’s solution was added on Day -1 to raise conductivity to the High Conductivity condition (300 ± 50 μS), which was stable 24h later (designated as Day 0). Measurements then continued from days 0 through 6 in the High Conductivity condition. The Control Condition (respirometry experiments only) followed the same experimental timeline, but fish remained in the Low Conductivity condition throughout the experiment.

### EOD Measurements

The EODs of seven *B. gauderio* and seven *E. virescens* were measured continuously during experimental days -2 through 6. Fish had constant access to food throughout the experiment.

Procedures for measuring calibrated EODs in free-swimming fish followed standard methods reported earlier (Stoddard et al., 2003). The measurement tank consisted of a 285-liter glass aquarium (120 cm x 44 cm x 44 cm) that was electrically shielded with grounded aluminum mesh screen. The aquarium was divided into three compartments with fiberglass screen panels and a mesh tube in the center compartment connected the outer two compartments such that the fish could swim between the outer compartments by passing through the center tube. A custom-built amplifier detected when the fish was centered in the tube, which then triggered a real-time digital processor (Tucker-Davis Technologies RP8, Gainesville, FL) to digitize the EOD at 48 kHz across a different pair of nichrome wires at opposite ends of the tank. EODs were amplified at 500x gain and low-pass filtered at 500 kHz (Cygnus FLA-01, Cygnus Tech, Delaware Water Gap, PA).

### Respirometry

The mass specific respiration rate (mM O2/g/min) of 14 *E. virescens* and 14 *B. gauderio* was measured once during the Low conductivity condition (day -1) and twice in the High conductivity condition (days 1, and 6). Under the Control condition, respiration was measured in additional *E. virescens* (n=14) and *B. gauderio* (n=14) following the same schedule as the experimental group (days -1,1, and 6).

Fish were acclimated to individual open flow respiration chambers for a minimum of 18 hours before respiration measurements to ensure fish were in a post-absorptive state so digestion would not affect their metabolic rate. The prolonged acclimation period also served to prevent transient changes in respiration rate caused by initial stress when fish are first placed in the chamber. Each fish’s length was recorded once at the start of the experiment and weight was recorded after each respiration measurement. EOD frequency (Hz) was recorded for the *E. virescens*, once, at the start of the experiment. To control for background oxygen consumption caused by the chamber’s microbial activity, background respiration was recorded immediately after each fish’s respirometry measurement from the empty chamber after the fish was removed, and respiration measurements were corrected according to recommended practices (Svenson et al., 2016).

The respiration chamber was a translucent acrylic cylinder, 24 cm in length and 5 cm in diameter, with threaded polyoxymethylene caps on each end (AlphaCool, Model 15719, Braunschweig, Germany). During acclimation, end caps had three 1-cm holes that were large enough to allow water exchange but small enough that fish could not swim out of the chamber. During respiration measurements, end caps with a single G1/4-threaded interface attached to quick-connect fittings (Colder Products Company, Roseville, MN) attached to aquarium tubing and a 9-volt aquarium pump.

During open circulation, oxygen saturated water from a large outer tank was pumped through the respiration chamber, while during closed circulation the pump circulated water through the respiration chamber without introducing new water from the outer tank. Open and closed circulation was controlled by connecting and disconnecting the tubing from the inflow side of the pump. The outflow side of the pump passed water through a custom-constructed acrylic plastic measurement chamber, where water oxygen concentration was measured with a NeoFox phase fluorometer (Ocean Optics, Orlando, FL), from a RedEye oxygen indicator patch (Ocean Optics) adhered to the interior of the measurement chamber. The RedEye oxygen indicator patch was calibrated using a two-point calibration method with the first calibration point taken at a dissolved oxygen level of zero, achieved by bubbling nitrogen into aquarium water, and the second point taken from aquarium water at full atmospheric oxygen saturation. Oxygen concentration was recorded from the NeoFox unit on a laptop PC using NeoFox Viewer software (Ocean Optics).

### Data Analysis

#### EOD Waveform Parameters

For both *B. gauderio* (n=7) and E *virescens* (n=7), EODa was measured from the negative minimum to the positive maximum of the EOD waveform (Figure 1). Additional EOD parameters for *B gauderio* included the amplitude of the EOD’s positive first phase (P1a) and the amplitude of the second negative phase (P2a), measured from zero volts to the positive peak or negative minimum, respectively. The durations of P1 and P2 were measured as the duration of each phase at 50% of peak amplitude (Figure 1).

**Figure 1.**
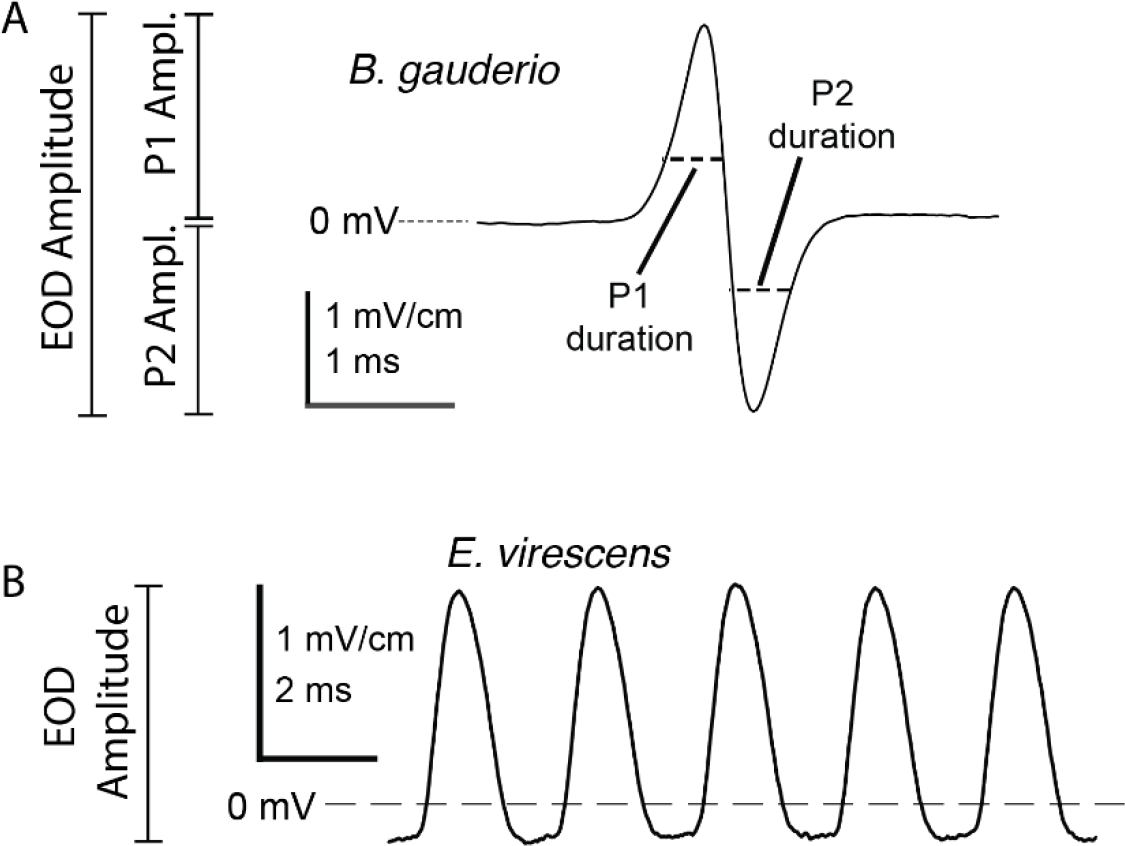
Measurement of EOD waveform parameters. (A) The EOD of *B. gauderio* is a biphasic pulse with an initial positive phase (P1) followed by a negative second phase (P2). The EOD amplitude (EODa) is measured from the positive peak to the negative peak, while P1 amplitude is measured from 0 mV to the positive peak, and P2 amplitude is measured from 0 mV to the negative peak. The durations of P1 and P2 are measured as the width of the phase at 50% amplitude. (B) The EOD of *E. virescens* is a monophasic positive pulse, repeated at high frequencies. Amplitude is measured from the waveform minimum to the positive peak.

Change in EOD waveform parameters over time and differences between species were analyzed with univariate analysis of variance (ANOVA). Comparisons between before and after rapid conductivity change assessed the immediate effects of increased water conductivity on EOD parameters. Comparisons between Day 0 and Day 6 assessed changes in the same EOD characters during continued exposure to High Conductivity. All analyses were performed with MATLAB (MathWorks, Inc. Natick MA). Data are presented as mean ± SEM.

#### Respirometry Analyses

Respiration rate was derived from a linear regression fit to the decline in water oxygen concentration during the closed circulation periods. Data were considered valid only if r^2^ for the regression exceeded 0.99. The corrections for background respiration and mass of each fish were calculated according to accepted standard practices (Svendsen et al., 2016). A two-way analysis of variance (ANOVA) was then used to compare the mass specific respiration rates across species and across water conductivities. Significant omnibus tests were further analyzed by post hoc pairwise comparisons with experiment-wise alpha maintained at 0.05 by Tukey’s HSD. A linear regression was also used to test for correlations of respiration rate with body condition index (BCI; weight (g) divided by length (cm)) in both species and a separate linear regression tested for a correlation of BCI with EOD frequency in *E. virescens*.

## Results

### Effects of Water Conductivity on EOD Waveform Parameters

In the Low Conductivity condition EODa was higher at baseline for *B. gauderio* (3.21 ± 0.48 mV/cm) than for *E. virescens* (1.14 ± 0.31 mV/cm; F_(1,12)_ = 13.03, p = 0.0036) (Figure 2). After the addition of Walter’s Solution on Day -1, water conductivity stabilized by mid-day on Day 0. The increased water conductivity resulted in decreased EODa from 3.21 ± 0.45mV/cm to 1.7 ± 0.30 mV/cm for *B. gauderio*, and from 1.15 ± 0.29 mV/cm to 0.89 ± 0.18 mV/cm for *E. virescens* (Figure 2) (F_(1,24)_ = 17.7, p = 0.003). After this initial decline, EODa increased over the course of 6 days in *B. gauderio* by 20.2% ± 4.3% but did not increase in *E. virescens*, changing by only -0.05% ± 6.1% (Figure 3) (F_(1,12)_ = 7.43, p = 0.018).

**Figure 2.**
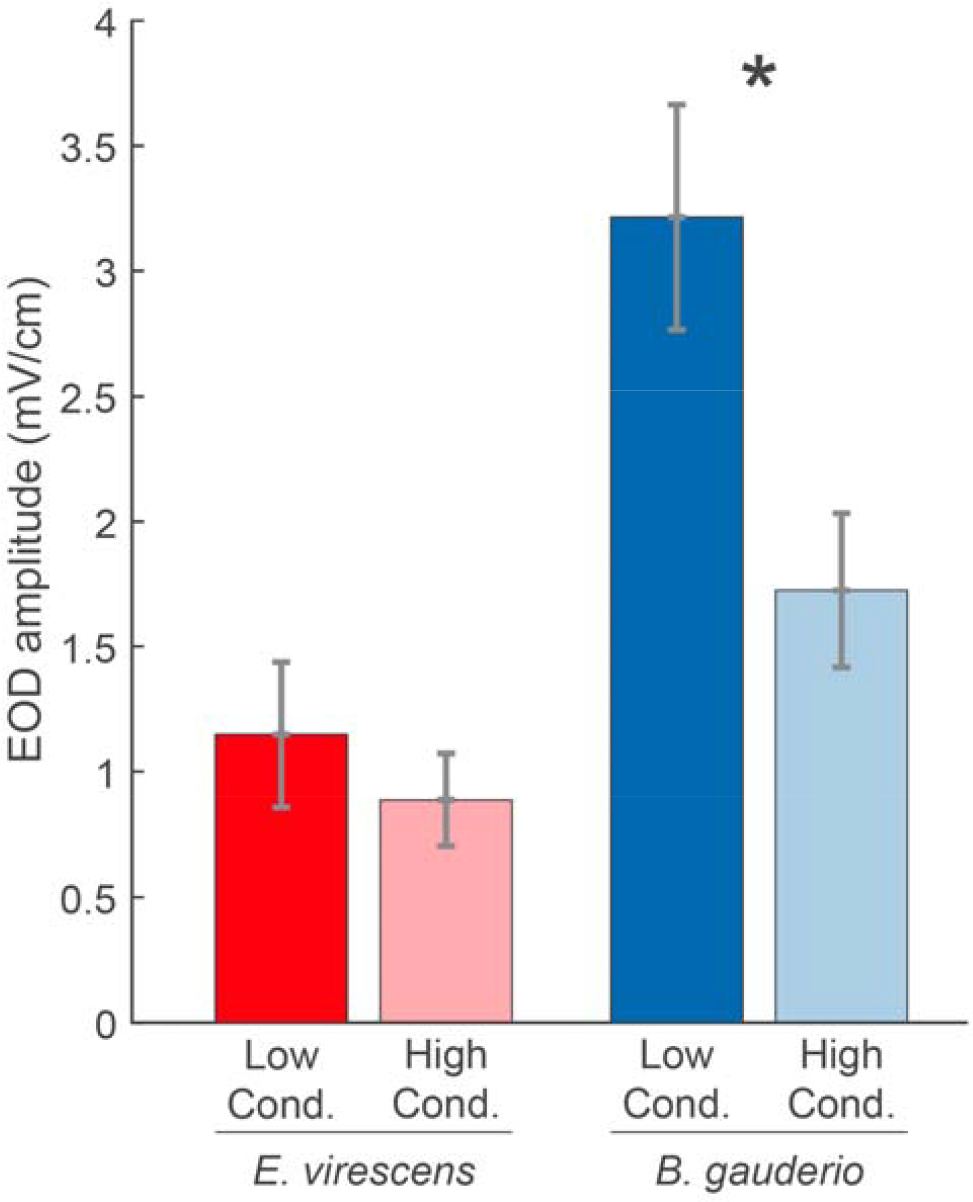
Change in EODa from low conductivity (dark bars) to high conductivity (light bars) for *E. virescens* (wave type; n=7) and *B. gauderio* (pulse type; n=7). While EODa decreased for both species, *B. gauderio*’s EODa had a much steeper decline in response to increased water conductivity than that in *E. virescens*.

**Figure 3.**
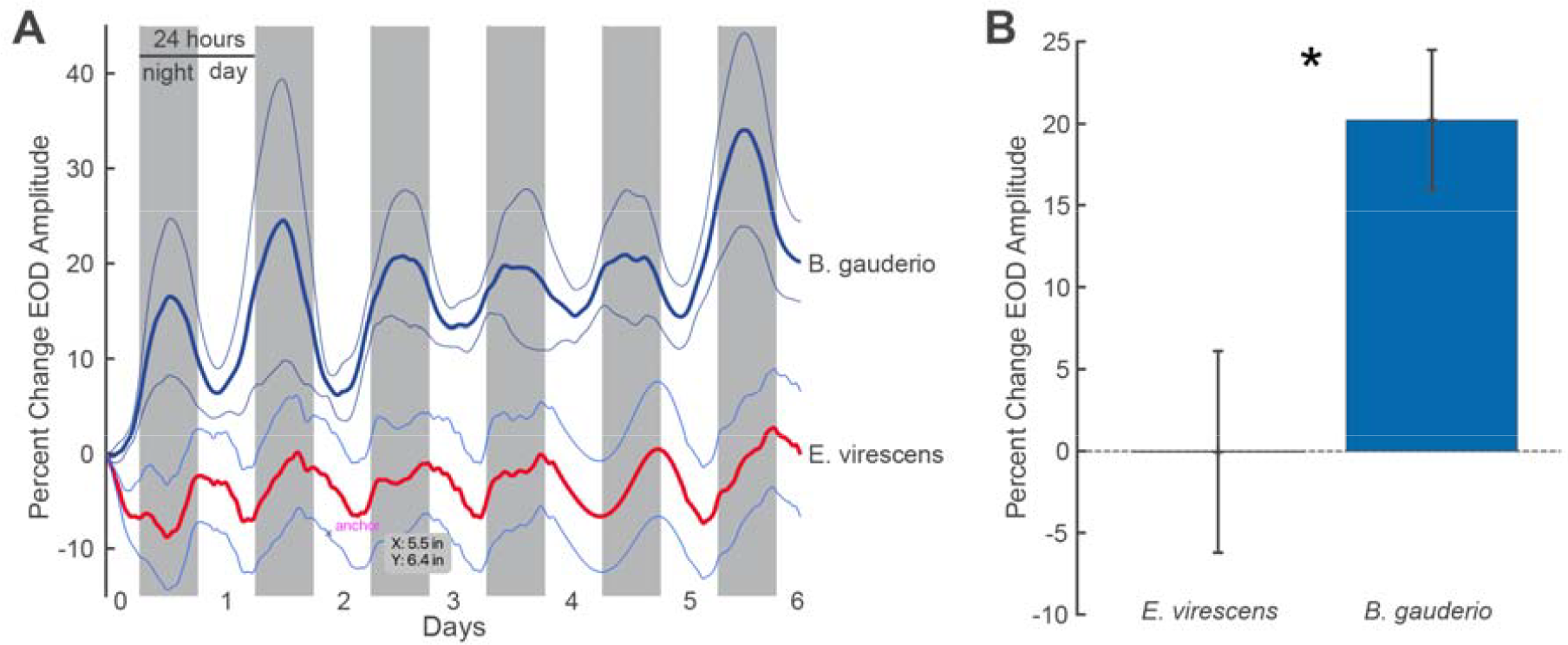
EOD responses of *B. gauderio* and *E. virescens* to increased water conductivity over six days. A) EOD amplitude (EODa) recorded for six days after water conductivity was increased from low conductivity (150 ± 50 μS) to high conductivity (350 ± 50 μS). Walter’s Solution was added on Day -1 to increase water conductivity, which was stable 24h later (designated as Day 0). Bold lines represent EODa normalized to mid-day on Day 0, while the thin lines indicate SEM. *B. gauderio* (n = 7) increased EODa gradually during the subsequent six days in high conductivity, but for the wave-type fish *E. virescens* (n = 7) EODa remained relatively constant. B) Percent change in EODa after six days in high conductivity water (D0 to D6). *B. gauderio* increased EODa by 20.2% ± 4.3% while *E. virescens* showed no major change in EODa (−0.05% ± 6.1%).

In *B. gauderio* EODa, P1a, and P2a parameters all increased in tandem during the 6 days in high conductivity, with P1a increasing by 21.8% ± 3.8% and P2a increasing by 26.6% ± 6.3% and no differences between EODa, P1a, and P2a after six days (Figure 4). The durations of P1 and P2 did not increase as did the EOD amplitude parameters, with P1 duration decreasing slightly by -1.3% ± 1.4%, and P2 duration decreasing by -0.3% ± 1.8% (Figure 4)(F_(4,30)_ = 11.14, p < 0.0001; post-hoc pairwise comparisons via Tukey’s HSD).

**Figure 4.**
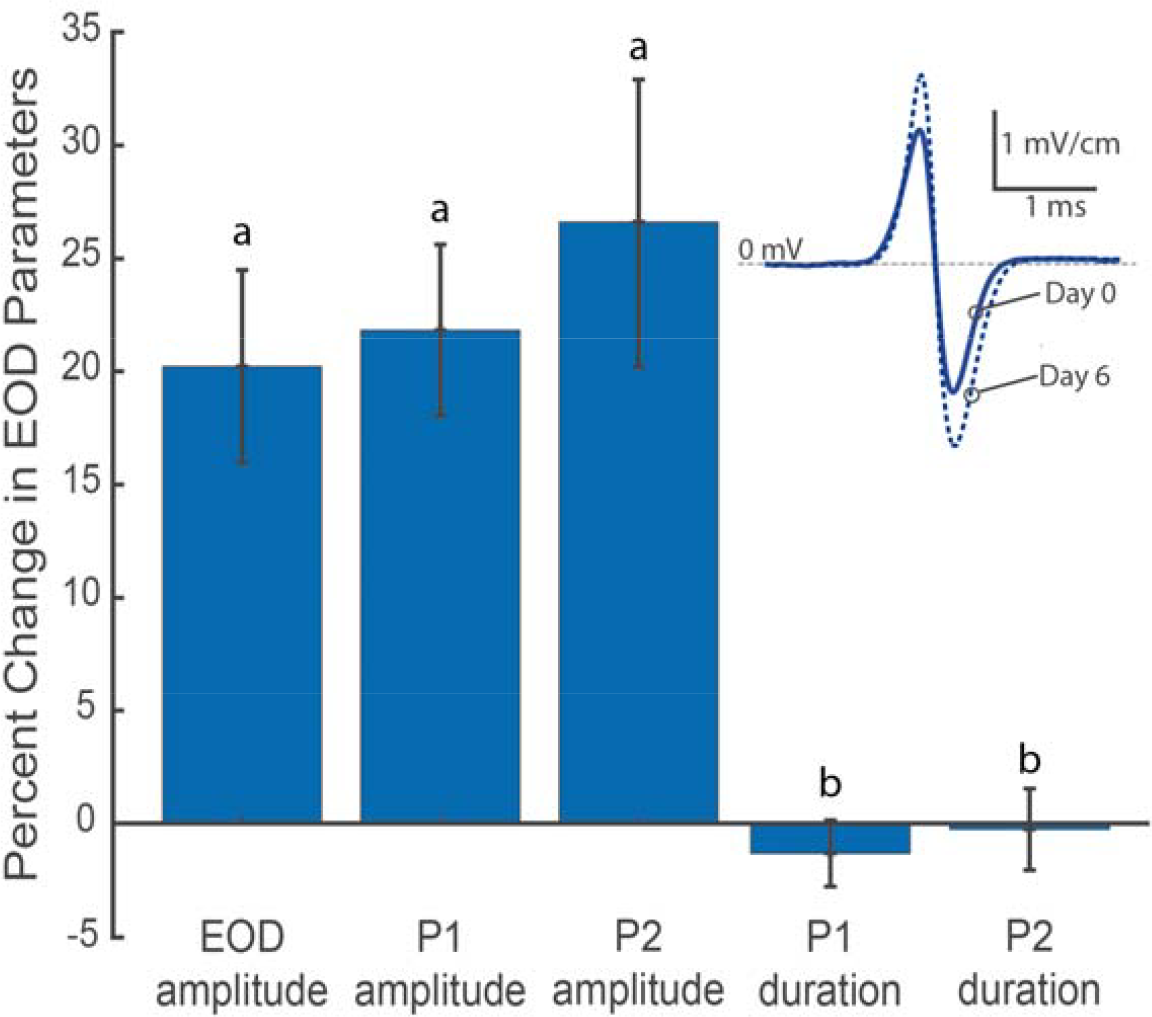
Percent change in EOD amplitude and duration parameters of the pulse-type *B. gauderio* (n=7) after six days in the High Conductivity water condition. EODs were recorded for six days following a water conductivity increase from low conductivity (150 ± 50 μS) to high conductivity (350 ± 50 μS). EOD amplitude increased in tandem with P1 amplitude and P2 amplitude, whereas P1 duration and P2 duration did not change. Parameters with the same lower case letter (a or b) are not statistically different, parameters with different lower case letters are statistically different (F_(4,30)_ = 11.14, p < 0.0001; post-hoc pairwise comparisons via Tukey’s HSD). **Inset** shows representative change in the EOD of a single *B. gauderio* from Day 0 (solid blue line) to Day 6 (dashed blue line) in high conductivity. On Day 0, water conductivity was increased from a low conductivity (150 ± 50 μS) to a high conductivity (350 ± 50 μS). The EOD P1 and P2 amplitudes increased in equal magnitudes from day 0 to day 6 while the EOD P1 and P2 durations remained constant.

### Effects of Water Conductivity on Metabolic Rate

Resting respiration rate was not correlated with BCI in *E. virescens* (R^2^ = 0.054, p = 0.39), whereas resting respiration rate had a weak positive correlation with BCI in *B. gauderio* (Figure 5; R^2^ = 0.198, p = 0.04). Because respiration rate was mildly correlated with BCI in only one species, we did not control for BCI during subsequent analysis.

**Figure 5.**
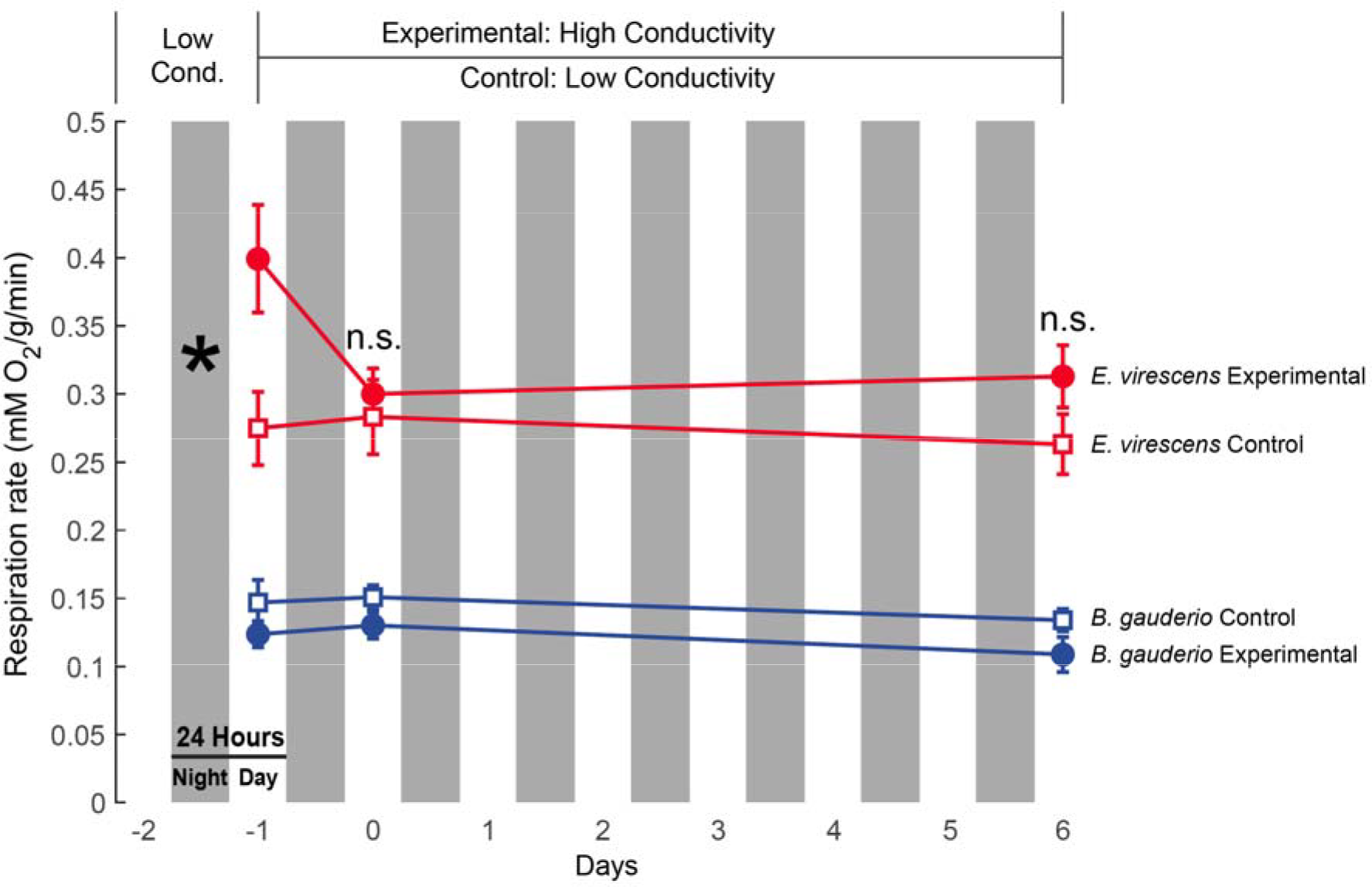
Metabolic responses of *B. gauderio* (n=14) and *E. virescens* (n=14) to increased water conductivity. The experimental values (circles) and control values (squares) show the respiration rates (mMO2/g/min) of in *E. virescens* (red) and *B. gauderio* (blue). Experimental treatment: Water conductivity increased from Low Conductivity condition (150 ± 50 μS) to a High Conductivity condition (350 ± 50 μS). Respiration was measured once in the low condition immediately before water was increased and then measured again on day 1 of the high condition and on day 6 of the high condition. Control treatment: water conductivity was kept at (150 ± 50 μS) and respiration rates were measured at the same intervals as the experimental treatment. The respiration rate of *E. virescens* decreased in high conductivity while the respiration rate of *B. gauderio* remained constant in both conditions. In all conditions *E. virescens* had a significantly higher respiration rate than *B. gauderio*. In the control condition, respiration rates for both species remained constant.

In all experimental conditions, the respiration rate was higher for *E. virescens* than for *B. gauderio* (Figure 5; species main effect: F_(1,175)_ = 141.4, p < 0.0001). After the addition of Walter’s Solution to increase water conductivity on Day -1, water conductivity stabilized by mid-day on Day 0. A significant species by conductivity interaction (F_(3,175)_ = 4.93, p = 0.0025), with post-hoc pairwise comparisons by Tukey’s HSD, supports that respiration rate in *E. virescens* decreased immediately after the change to the High Conductivity (p=0.0034) condition while the respiration rate of *B. gauderio* remained constant after the change to High Conductivity (p=1.0000). The respiration rates for both *E. virescens* and *B. gauderio* did not change over six days in High Conductivity. In the Control condition, respiration rates for both species remained constant throughout the experiment.

## Discussion

### Species Differences in Response to Increased Water Conductivity

Following a rapid increase in conductivity, EODa decreased in accordance with Ohm’s law only in the pulse-type *B. gauderio*, while the wave-type *E. virescens* unexpectedly showed only a small decrease in EODa. During the following six days of stable high conductivity, *B. gauderio* began recovering EODa within 48 hours, ultimately increasing EODa by approximately 20% after 6 days. Surprisingly, during the rapid conductivity increase and the six subsequent days in high conductivity, there was no change in metabolic rate for *B. gauderio*, even as EODa increased, while metabolic rate decreased after the rapid increase in water conductivity for *E. virescens* even as EODa remained constant.

Since EODa determines the fish’s electrosensory range (Assad et al., 1999; Nelson & MacIver., 1999), these results suggest that *B. gauderio* prioritizes the maintenance of sensory range during periods of increased water conductivity through some mechanism of recovering EODa that does not increase total metabolic rate. Under the same conditions, the EODa of *E. virescens* did not decrease as expected when measured several hours after the addition of a saline solution to the water, suggesting perhaps a rapid physiological mechanism that adjusts EODa to changes in conductivity faster than our measurements could detect. The decreased metabolic rate observed in *E. virescens* may or may not be associated with any such mechanism. Several factors, alone or in concert, could account for the widely divergent responses of these species to disruptions in electrosensory and electric communication performance after sudden increase in water conductivity.

### Habitat and Reproductive Life History of B. gauderio *and* E. virescens

Differences in habitat offer one potential explanation for the observed differences between species. *E. virescens* inhabits deep waters with historically stable water conditions (Silva et al., 2003), whereas *B. gauderio* inhabits a variety of aquatic environments including riverbanks, slow-moving creeks, and floodplains (Giora & Malabaraba, 2009). Thus, *B. gauderio* may be adapted to habitats where rapid changes in water conductivity are common, whereas *E. virescens* may be suited for less variability than *B. gauderio* and is instead well-adapted for a limited range of change.

Divergent reproductive life histories could also explain why *B. gauderio* restores EODa while *E. virescens* does not. Semelparous species, such as *B. gauderio*, have one breeding period before death while iteroparous species, such as *E. virescens*, have multiple breeding periods over the species’ lifespan. Urgency to mate, such as a terminal investment in reproduction, could in part explain why *B. gauderio* responds to increased conductivity by directing effort into EODa recovery, thus prioritizing communication. A similar pattern of preserving communication during stress occurs when *B. gauderio* males increase both EODa and EOD duration in response to food deprivation, again perhaps as a terminal investment in reproduction (Gavassa & Stoddard, 2012).

In contrast, the decrease in the metabolic rate of *E. virescens* could be a characteristic of iteroparity, possibly suggesting an effort to conserve energy and wait for conditions to improve. However, this is inconsistent with earlier reports of that behavior where *E. virescens* reduced energetic costs under hypoxia by diminishing EODa, but without an associated reduction in metabolic rate (Reardon et al., 2011). Furthermore, when under metabolic stress from food deprivation, *E. virescens* reduces EODa over several days, possibly as a reproductive strategy to conserve energy until conditions improve (Sinnett & Markham, 2015).

### Mechanisms for EODa Compensation in B. gauderio

Contrary to our expectations, *B. gauderio* showed no increase in metabolic rate associated with their gradual increase in EODa. This outcome can only be explained in one of two ways: either *B. gauderio* recovers EODa by a mechanism that requires no additional metabolic investment or by increasing metabolic investment in EO output while reducing metabolic investment in other physiological processes.

In all studies published to date where *B. gauderio* increases EODa by increasing electrocyte power output and metabolic investment in EOD production, increased EODa is accompanied by corresponding increases in EOD duration, P1 duration, and P2 duration (Markham & Stoddard, 2005, 2013; Salazar & Stoddard, 2008). This was not the case in the present study, wherein *B. gauderio* increased EODa with no accompanying changes in EOD duration, P1 duration, and P2 duration, suggesting a different physiological mechanism is at work. If *B. gauderio* is instead increasing EODa without additional metabolic investment in EO output, perhaps the only feasible mechanism for doing so is impedance matching. When increased water conductivity reduces the impedance of the water (the load impedance) the fish may be reducing the internal resistance of the EO (the source impedance) to maximize power transfer from the EO into the water in the new conductivity condition. One mechanism for reducing the internal resistance of the EO would be to decrease the electrocyte membrane resistance, which is known to occur in response to stress and social activity in other gymnotiforms (Markham et al., 2009), thereby reducing the internal resistance of the EO without requiring additional metabolic investment.

The second possibility is that *B. gauderio* invests more energy in EOD production by diverting metabolic resources from other physiological processes. These types of metabolic tradeoffs are common among animals (Stearns, 1989; Zera & Harshman, 2001; Moore & Hopkins, 2009). For example, when exposed to increased water temperatures, the resting metabolic rate (RMR) of the Antarctic fish, *Trematomus bernacchii*, increases initially in a temperature-dependent fashion. The RMR then returns to baseline levels over the course of 9 weeks but that happens with an associated decrease in body mass (Sandersfeld et al., 2015). Another example can be seen in the trade-off between energy conservation and increased predation in the mourning dove, *Aristolochia macroura*. During the winter months when food availability is low, *A. macroura* uses regulated nocturnal hypothermia to conserve energy. However, this energy saving behavior comes at the expense of increased predation due to an associated reduction in flight ability (Carr & Lima, 2013).

### EODa Stability in E. virescens

In accordance with Ohm’s law, EODa should be reduced by half when water conductivity is increased by a factor of two but surprisingly, for *E. virescens*, EODa decreased very little after water conductivity was doubled. In this experiment, the effects of conductivity on EODa were assessed by comparing EODa before addition of the saline solution to EODa measured after conductivity had stabilized several hours later. Measured in this manner, the EODa of *E. virescens* did not decrease significantly. One possible explanation is that *E. virescens* employs a rapid mechanism to compensate for changes in conductivity within a matter of minutes or hours. Such changes would not have been detected with the methodology used in the present study.

### E. virescens *Reduces Metabolic Rate When Conductivity Increases*

Another surprising finding of this study was that *E. virescens* showed a large decrease in metabolic rate after water conductivity increased. One possible explanation is that this decrease in metabolic rate is the result of lower activity levels in electrosensory areas of the brain. Electrosensory processing in the brain incurs high metabolic costs (Sukhum et al., 2016; Nilsson, 1996), driven in part by large populations of neurons firing 1:1 with the EOD rate (Salazar et al., 2013). Reduced EODa in high water conductivity could reduce activity levels in these neuronal populations, thereby reducing overall metabolic demand in peripheral and central neural systems that encode and process electrosensory information (Sukhum et al, 2016). However, because the decrease in EODa was not significant for *E. virescens*, further investigation is needed to test this possibility.

An important limitation of this work is that we exposed fish to only two different water conductivities. Perhaps the shift from 150 ± 50 μS to 350 ± 50 μS was not large enough to elicit a metabolic response from *B. gauderio* or an EODa response in *E. virescens*. Another variable not considered within the study design was the normal water conditions in each species’ habitat. Instead, the conductivities were standardized for both species creating the possibility that one or both conductivity phases could have unintentionally been more favorable to one species. Sex differences, age, and the developmental stage of each fish were not considered due to limitations in fish available. These variables should be explored in future studies.

Further research is needed to address how weakly electric fish respond to anthropogenic disturbances in water quality and should include a larger diversity of species and a wider range of water conditions. This includes examining response differences between species who differ significantly in EOD frequencies, species whose EO is derived from muscle tissue (myogenic) versus neural tissue (neurogenic), and comparisons between Neotropical gymnotiforms and Afrotropical mormyrid electric fish. Further investigation involving different magnitudes of change in water conductivity is also needed to determine how *E. virescens* responses to increased water conductivity. Additionally, the effects of rapid transitions from high to low conductivity (opposite to this study), and prolonged exposure to altered water conductivity should be explored.

Continued examination is necessary to determine what sensory mechanism in *B. gauderio* initiates the increase of EODa in high conductivity. Possible mechanisms include the direct sensing of water osmolarity or conductivity, or electrosensory detection of the reduction in EODa. It will be important to assess the nature and extent of how changes in EODa affect sensory performance with respect to navigation, object detection, and prey detection/capture. Additionally, future research should examine whether communication or social interactions are altered or impaired after increases in water conductivity.

## Conclusion

The negative impacts of anthropogenic activity on South American freshwater ecosystems are expected to accelerate (IPCC, 2022) if preventative actions are not immediately implemented (Kuemmerlen et al., 2022). As a result, electric fishes may be disproportionately harmed relative to other freshwater fishes as pollution harms physiology in general, but also potentially degrades their primary sensory and communication modalities (Markham et al., 2016). The findings of this study suggest that weakly electric fish may display widely divergent responses to changes in water conditions and emphasizes that a more comprehensive and comparative analysis of how electric fish respond to disturbances in aquatic habitats will be essential for predicting and perhaps mitigating the consequences of anthropogenic activity to neotropical aquatic habitats.

## Acknowledgements

We would like to thank Rosalie Maltby for laboratory support and fish care. We are grateful to Caryn Vaughn and Ricardo Betancur-R. for comments on an earlier version of this manuscript. This data presented in this manuscript were reported in a master’s thesis submitted by S.D.W. to the University of Oklahoma, available at this URI: https://shareok.org/handle/11244/336889.

## Data Availability

All data are available immediately upon request to the corresponding author.

